# Biological tick control: modeling the potential impact of entomopathogenic fungi on the transmission of East Coast fever in cattle

**DOI:** 10.1101/2024.07.08.602534

**Authors:** Joseph Wang’ang’a Oundo, Nienke Hartemink, Mart C.M de Jong, Constantianus J.M. Koenraadt, Shewit Kalayou, Daniel Masiga, Quirine ten Bosch

## Abstract

Biological control of ticks using entomopathogenic fungi (EPF) is a highly desired alternative to chemical acaricides for the control of tick-borne pathogens. For *Metarhizium anisopliae* isolate ICIPE 7, one of these EPFs, efficacy against multiple tick species has been demonstrated in laboratory and field settings. However, we currently have little quantitative understanding of how EPFs can impact transmission. We developed a deterministic model of tick–host–pathogen interactions to explore how the effects of EPF on *Rhipicephalus appendiculatus* ticks may impact the transmission dynamics of East Coast fever (ECF) in cattle populations. We parameterized the multi-faceted effects of EPFs on tick dynamics using experimental data on Tickoff® biopesticide (a novel formulation of *M. anisopliae* ICIPE 7) and related EPFs. The epidemiological impact of EPF was evaluated across a range of product profiles and implementation strategies. Model results indicate that, for the explored product profiles, EPF derives most of its epidemiological impact through the delayed mortality effect. This EPF-induced mortality could not only reduce the onward *Theileria parva* transmission to cattle (both treated and untreated) but could also cause a reduction in the tick-to-host ratio and thus cattle exposure to ticks. The effects of EPF on reproduction fitness and engorgement of ticks elicit negligible impact. High levels of population coverage and treatment frequency are needed to reduce the tick population size and reach meaningful epidemiological impact in cattle populations. Additionally, increasing the persistence time of fungal conidia on cattle skin – through technological improvements to the EPF formulation – can substantially reduce acute infections when combined with appreciable population coverage levels, treatment frequency, and efficient spraying techniques. Our model analysis provides insights into the potential impact of EPF when deployed at a population level, and lends support to further research and development of this biological tick control tool.

## Introduction

East Coast fever (ECF) disease is a major constraint to cattle health and productivity in 11 countries in eastern, central, and southern Africa, including Kenya (Gachohi et al., 2012; Nene et al., 2016). The disease is caused by the protozoan parasite *Theileria parva* and is transmitted by the three-host ixodid tick *Rhipicephalus appendiculatus*. The African Cape buffalo (*Syncerus caffer*) is the natural reservoir host for *T. parva* (Nene et al., 2016). The economic impact of ECF includes reduced meat and milk production, cattle morbidity and mortality, and control measure costs against both ticks and the disease (Gachohi et al., 2012; Nene et al., 2016). These economic losses tend to affect resource-poor households disproportionately. In recent years, the geographic range of *T. parva* has expanded, for example to non-endemic countries of Comoros island (De Deken et al., 2007) and Cameroon (Silatsa et al., 2020).

Control of tick-borne diseases in cattle has traditionally relied on the use of chemical acaricides to kill the tick vector. The effectiveness of chemical acaricides for the control of tick infestation in cattle has been demonstrated in several field trials (Muraguri et al., 2003; Murigu et al., 2016; Nonga et al., 2012). However, the long-term sustainability of this tick control method is threatened by the emergence of acaricide resistance in ticks, including *R. appendiculatus* ticks (Githaka et al., 2022; Ntondini et al., 2008; Vudriko et al., 2016), and concerns regarding contamination of the environment and milk and meat products (De Meneghi et al., 2016). The intensive and frequent use, as well as inappropriate use of these chemicals, has been associated with the development of acaricide resistance in ticks (Githaka et al., 2022; Vudriko et al., 2018). There is, therefore, a need for new and environmentally friendly alternatives for tick control.

Biological control of ticks using entomopathogenic fungi (EPF), especially *Metarhizium anisopliae* sensu lato (s.l.) and *Beauveria bassiana* s.l., has attracted much interest as a possible and valuable alternative to conventional chemical acaricides. A range of laboratory studies has demonstrated the ability of EPFs to cause high mortality in the larva, nymph, and adult stages of various tick species (Hedimbi et al., 2011; Kaaya et al., 1996; Kaaya and Hedimbi, 2012). However, the success of tick control under field conditions has had variable results, in that, some studies reported a substantial efficacy (Alonso-Díaz et al., 2007; Barbieri et al., 2023; Murigu et al., 2016) while others reported a lack of significant efficacy when compared to the respective controls (Correia et al., 1998; Leemon et al., 2008; Oundo et al., 2024). The success of EPFs in the field is influenced by environmental factors such as temperature, relative humidity, and solar ultraviolet (UV) radiation (Fernandes et al., 2012). Therefore, this discrepancy in the outcomes underscores the need for further research and development of EPFs before their adoption can be recommended and implemented at a wider scale.

Beyond increasing the mortality rate of ticks, EPFs also elicit a multitude of effects on the infected ticks including a reduction in engorgement weight, fecundity (egg mass weight), and egg hatchability. Besides, increases in periods of engorgement, preoviposition, oviposition, and post-oviposition have also been observed (Nana et al., 2015, 2012). However, we currently lack a comprehensive quantitative understanding of how these effects interact to impact ECF transmission in cattle. While EPFs do not cause instantaneous tick mortality, the hallmark of chemical acaricides, they will still kill the infected ticks before they can molt to become infectious and thereof transmit the infection to the next host. This slow mortality rate implies that EPFs have limited potential to provide direct protection from infective tick bites at the individual animal level, but may still offer community-level protection. However, the level of coverage within the population, the duration of persistence on treated cattle, and the frequency and duration of treatment application required to achieve a maximum impact are unknown. Mathematical models can help to improve our quantitative understanding of how EPFs can indirectly protect the overall cattle population from tick-borne infections.

Mathematical models have been used to evaluate different control strategies for ECF in cattle populations. The study by Walker et al. (2014), for example, illustrated that the elimination of *T. parva* infection in cattle is unlikely to be accomplished solely by frequent acaricide use on cattle when grazing land is shared with the reservoir host Cape buffalo. This work builds on the earlier recognition by Medley et al. (1993) that the interruption of transmission of *T. parva* infection through tick control requires drastic reductions in tick infestations. Their modeling approaches did not incorporate the infection dynamics within the tick population and did not explicitly include the development stages (egg, larva, nymph, adult) of the tick vector. As the transmission cycle of ECF and other tick-borne pathogens encompasses several tick development stages, each of which may be affected differently by EPFs, these frameworks may not be suitable for investigating the multifaceted effects of EPFs.

Here, we introduce a detailed deterministic model of tick–pathogen–host interactions developed to estimate the impact of EPFs on the transmission of ECF in the cattle population managed under an extensive grazing system. We used the model to explore the implementation strategies and product properties needed to achieve a meaningful epidemiological impact. This model simulation is not intended for prediction of a specific product, but rather to provide an illustrative framework to improve our understanding of the potential benefits that can be accrued when EPF is deployed at a population level.

## Model development

### Study setting/system

We are simulating the impact of EPF on a cattle population managed under an extensive grazing system where cattle are allowed to graze on natural pasture on fallow or communal grazing land. There is no controlled rotational grazing and animals have access to the entire grazing area. Cattle in this grazing system are exposed to tick reinfestation from the environment throughout the year and hence are at a constant risk of tick-borne infections. The tick control practice consists mostly of regular biweekly treatments and the treatment coverage level is limited, reaching a maximum of 40% of the population. This study setting allows us to assess the practical impact of EPF within the context of this herd management system and tick control efforts.

We constructed a tick–pathogen–host interaction model, describing the transitions between infection and treatment states for cattle and the different life stages of ticks (Figure 1; Appendix A.1. Times to events are assumed to be exponentially distributed, so that the average duration of a state/stage is the reciprocal of the rate of leaving that state/stage. The ordinary differential equations (ODEs) describing the full model are presented in the supplementary material (Appendix A.2). In this continuous-time, stage-structured deterministic model, we describe the population dynamics of *R. appendiculatus* ticks and cattle, the transmission of *T. parva* within the tick vector and between the tick and host population, and the application and decay of the treatment.

**Figure 1.**
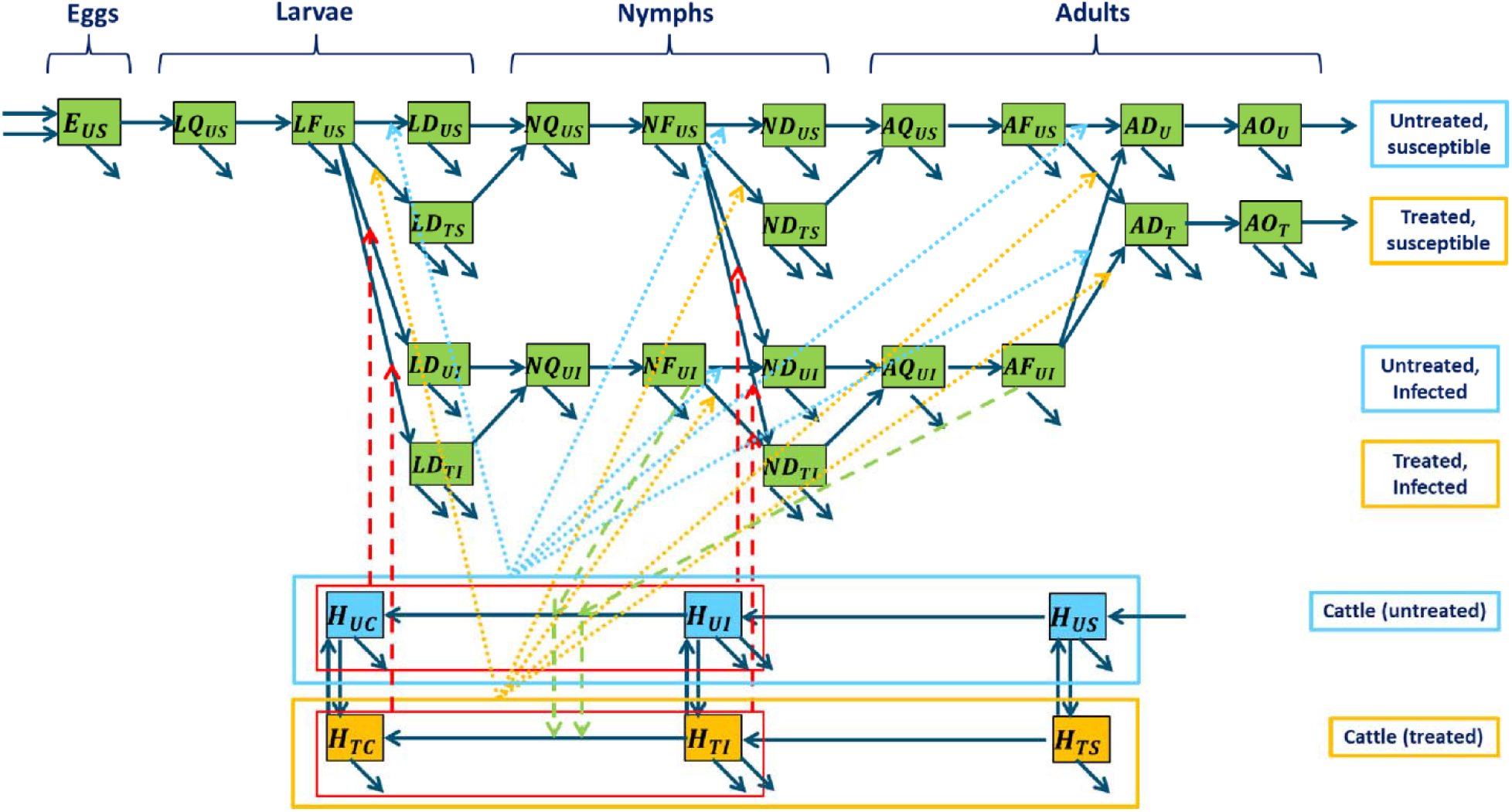
Conceptual model representing the tick–pathogen–host–treatment interactions. For the tick compartments, the capital letters indicate the developmental stage (E-egg, L-larva, N-nymph, A-adult) and the physiological phase (Q-questing, F-feeding, D-development, O-ovipositing). Host compartments are denoted with an H. Subscripts represent the treatment status (U – untreated or T – treated) and infection status (S – susceptible and I – infected/infectious, for hosts and ticks, and C – for carrier hosts). Solid navy-blue arrows denote demography, developmental, treatment, or pathogen-state transitions; green dashed arrows denote tick-to-host transmission routes; dashed red arrows denote host-to-tick pathogen transmission and/or biopesticide treatment; dotted arrows denote contact of the tick with EPF (light blue: no tick contact with the treatment, yellow: tick contact with the treatment).

### Tick and host population dynamics

The life cycle of *R. appendiculatus* involves four successive development stages, namely egg (E), larva (L), nymph (N), and adult (A). Except for the egg stage, each tick could either be in the questing (Q), feeding (F), or interstadial development (D) phase. Adult females can also be in the oviposition (O) phase. A female tick that survives to reproduce will consume three blood meals in its lifetime. Each of these blood meals will occur on a different host individual. In this model, we assume that the ticks feed only on cattle hosts. This assumption is based on the notion that *R. appendiculatus* is well adapted to the presence of domestic cattle and can be maintained by all stages feeding on cattle (Walker et al., 2003). In the study area of interest, wildlife such as buffaloes, elands, waterbucks, nyalas, greater kudus, and sable antelopes, which are alternative hosts for this tick (Walker et al., 2003), are not present.

After taking a blood meal from the cattle host, the female ticks will detach and find a suitable location in the environment to lay their eggs (*E*). A fully engorged adult female *R. appendiculatus* will lay 3,000 to 5,000 eggs and then die (Walker et al., 2003). The production of eggs is assumed to be proportional to the total number of ovipositing adult female ticks (*AO*_*U*_) and the egg-laying rate:

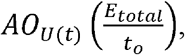

which is the reciprocal of the average time between each oviposition event. The eggs produced will hatch and develop into the questing larval stage at a constant developmental rate *k*_*E*_. Questing larvae will attach to cattle hosts at a constant rate *α*_*L*_ to feed. The feeding larvae will engorge with a mean duration *d*_*L*_ then detach from the host and enter the development phase. Larvae in the development phase will molt to the questing nymphal stage at a developmental rate *k*_*L*_. The same process of questing (*α*_*N*_, *α*_*A*_)feeding (*d*_*N*_, *d*_*A*_), and development (*k*_*N*,_ *k*_*o*_) is repeated for nymphs and adults. After this development phase, adult ticks mate and only the females proceed to the oviposition phase (through a sex proportion ζ). The ovipositing females will lay eggs for a duration *t*o and the depleted female will then die. The natural mortality rates are represented by μ_*ij*_ where *i* and *j* represents the tick developmental stages and phases respectively.

For the cattle, we assume a constant population size, which is obtained by keeping the birth rate and the death rate the same (*µ*_*H*_). Since there is no vertical transmission of the disease, all newborn cows are assumed to be susceptible and enter the *H*_*US*_ class.

### *Theileria parva* transmission dynamics

#### Host to tick transmission

*Theileria parva* infection is acquired from an infectious host by larvae or nymphs, maintained trans-stadially through the tick’s development and molting processes, and transmitted to a susceptible host by the next tick stage (nymph or adult). Infection acquired by adult ticks cannot be transmitted transovarially via the eggs to larvae of the next generation (Nene et al., 2016). Our model therefore ignores the infection acquired by the adult ticks. The model also assumes that ticks do not die of the *T. parva* infection and remain infectious for the remainder of their lives with 100% transstadial transmission. The probability of acquiring *T. parva* infection from the infectious cattle by larvae and nymph depends on whether it feeds on the acutely infectious host (*p*_*LI*_, *p*_*NI*_) or the persistent carrier host (*p*_*LC*_, *p*_*NC*_). The tick population is sequestered into a susceptible class and an infected/infectious class (second index subscripts: S – susceptible, I – infected/infectious).

#### Tick to host transmission

The total cattle host population (*H*_*total*_) is divided into sub-categories depending on treatment status (using subscript U for untreated or T for animals treated with EPF). This is further divided into susceptible (*H*_*US*_ and *H*_*TS*_), symptomatic infectious (*H*_*UI*_ and *H*_*TI*_), and carrier (*H*_*US*_ and *H*_*TC*_) compartments. Individuals move between compartments when their disease status and/or treatment status changes.

The susceptible host moves to the symptomatic infectious class after getting a bite from an infectious nymph or adult tick. The force of infection in the susceptible cattle (i.e., the rate at which the susceptible cattle become infected) is determined by the cattle exposure rate to ticks, the probability that the bite is by an infectious tick, and the probability of transmission per bite (*P*_*HN*_ and *P*_*HA*,_ for nymphs and adults, respectively). The cattle exposure rate to ticks, defined here as the average number of tick bites per host per day, is calculated as the ratio between the number of questing ticks and the number of hosts 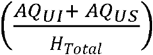 or 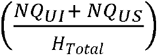 representing the number of questing adults or nymphs available per host, multiplied by the attachment rate of questing ticks (*α*_*A*_ and *α*_*N*_ for adults and nymphs).

For the probability that the bite is taken by an infectious tick, we use the proportion of feeding ticks that are infectious 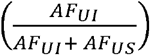 and 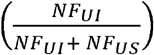, for adults and nymphs respectively.

The transmission probability per bite is *P*_*HN*_ for infectious nymphs and *P*_*HA*_ for infectious adults. A proportion (*P*_*I*_) of symptomatically infectious cattle may experience disease-induced death at a constant rate (*µ*_*I*_), while surviving individuals progress to the carrier compartment at the rate σ_*H*_.

We parameterized the model such that the probability of hosts dying from ECF is 5% (Medley et al., 1993). Cattle in this compartment develop solid immunity against re-infection with similar strains (Nene et al., 2016) and remain persistent carriers of tick-transmissible infection (Kariuki et al., 1995; Olds et al., 2018).

### Treatment with entomopathogenic fungi

Spraying of untreated cattle host (*H*_*US*_, *H*_*UI*_ and *H*_*UC*_) with EPF at the rate cp will produce a treated host population (*H*_*TS*_, *H*_*TI*_ and *H*_*TC*_). The treated host population also loses the treatment status over time, due to the decay of the conidial spores; we assume a constant decay rate (*δ*) (i.e. the rate of losing treatment status). In the absence of concrete data, we assumed a conservative estimate for *δ* of 1.0 per day. Thus, the EPF biopesticide is presumed to exhibit an effective acaricidal activity lasting one day, on average. Ticks attached to the cattle at the time of treatment will contact the EPF with varying probabilities for larvae (*P*_*LT*_), nymphs (*P*_*NT*_) and adult ticks (*P*_*AT*_). The on-host ticks that will contact the treatment will progress to treated status while those that escape treatment will remain in the untreated status (first index subscripts: U – untreated or T – treated with EPF).

### Effects of entomopathogenic fungi

The entomopathogenic fungi *Metarhizium anisopliae* ICIPE 7 elicits multifaceted effects on *R. appendiculatus* ticks, including increasing the overall tick mortality rate, prolonging the engorgement period, and interfering with the reproductive fitness of engorged female ticks by reducing fecundity (egg mass), and increasing pre-oviposition, oviposition and post-oviposition periods (Nana et al., 2015).

### Increased mortality

Unlike chemical acaricides which cause rapid death of exposed ticks, EPFs will take several days to kill a tick after exposure. We included this delayed lethality in our model by setting the EPF-induced mortality rate *ϑ*_iDF_ (where *i* = L, N, A) at the interstadial development phase instead of the feeding phase. The EPF-induced mortality *ϑ*_iDF_ was derived by fitting a Weibull model to tick mortality data (Oundo et al., 2024). The death rate increased by a factor of 10.

### Increased engorgement duration

In addition to increasing mortality, *M. anisopliae* ICIPE 7 may prolong the engorgement duration in treated *R. appendiculatus* ticks by 37.6% (Nana et al., 2015). We explore this in our model by increasing the engorgement duration (*d*_*i*_ where *i* = L, N, A) by *τ*_*i*_ days, resulting in an overall delay in detachment from the host. Thus, the rate of leaving the feeding compartment for treated ticks is assumed to be 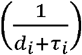.

### Decreasing reproduction fitness

Fungal infection can reduce the reproductive fitness of engorged females of *R. appendiculatus* ticks through a reduction in the fecundity rate by 36.9%, and an increase in the pre-oviposition, oviposition, and post-oviposition periods by 38.9%, 24.4% and 37.9% respectively (Nana et al., 2015). We explore this by increasing the pre-oviposition period (*k*_*o*_, i.e., the interval that elapses between the detachment of an engorged female and the first appearance of eggs) by *k*_*τ*_ days. Thus, the rate at which treated ticks leave the preoviposition phase is expressed as 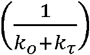. The fecundity (i.e., the average number of eggs laid per female tick) is reduced by a factor TFR (treatment fecundity reduction), while the oviposition duration is increased by τ_*o*_ days. This means that the rate of leaving the oviposition phase is assumed to be 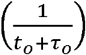, and the expression for the egg-laying rate becomes:

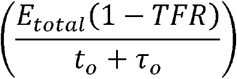

Where *E* is the number of eggs laid per ovipositing female, TFR is the treatment fecundity reduction, *t*_*o*_ is the normal oviposition duration in untreated adult females, and *τ*_*o*_ is the increased oviposition duration.

### Parameterization

The tick model is based on the assumption that, without treatment, the *R. appendiculatus* population is at equilibrium, that is, there is neither exponential growth nor decline of the tick population over time. We calibrated the model to achieve an equilibrium state where one egg-laying female tick replaces herself per generation (Randolph, 2004, 1998). If each female lays approximately 3000 eggs, the tick population equilibrium requires 3.9% survival from eggs to larvae, 9% survival from larvae to nymphs, and 19% survival from nymphs to fully reproductive adults (Randolph, 1998). We also calibrated the model such that the lifecycle duration of one generation of tick lasts for 276 days as observed in a previous field observation study (Branagan, 1973a). Besides, the equilibrium prevalence of *T. parva* in the cattle population has been calibrated to range between 7-10% as observed in our earlier study (Oundo et al., 2022).

The growth rate of the tick population within an ecosystem is assumed to be density-dependent, meaning the tick population growth rate increases when the population is low but slows down when the population approaches the carrying capacity. This density-dependent growth rate of the tick population follows a characteristic logistic model depending on the environment’s carrying Capacity *K*_*T*_ (i.e., the maximum number of ticks an environment can sustain for an indefinite period given resource availability), and is expressed as:

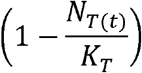

Where:

*N*_*T*_ – is the total tick population size

*K*_*T*_ – environmental carrying capacity for the tick population

The carrying capacity for the environment was set to be seven times larger than the total tick population at the equilibrium state.

### Model simulations of implementation strategy and product profile

The model simulations were implemented using the package ‘deSolve’ (Soetaert et al., 2010) in R version 4.3.1. Parameter values used to simulate the model are summarized in Table 1. We evaluated the potential impact of EPF by simulating different implementation strategies and product properties: (1) the application of biopesticides to cattle population at varying coverage levels (defined here as the proportion of cattle within the population that is treated with EPF) and treatment intervals for one year, (2) efficiency of treatment application (spraying) technique, (3) the duration of persistence of the EPF on cattle skin post-treatment, and (4) the different combinations of the effects of *M. anisopliae* isolate ICIPE 7 on the tick life cycle. Simulations (1) and (2) are implementation strategies while (3) and (4) are product properties. The impact of EPF was assessed based on the number of susceptible cattle that get infected.

**Table 1.** Parameter estimates used in the model.

| Parameter | Description | Value | Unit | References |
| --- | --- | --- | --- | --- |
| $E$ | Maximum number of eggs laid per ovipositing female | 3500 | eggs | (Walker et al., 2003) |
| $k_o$ | Preoviposition period of the engorged female tick | 6 | days | (Branagan, 1973b) |
| $t_o$ | Oviposition period of the engorged female tick | 24 | days | (Branagan, 1973b) |
| $k_E$ | Interstadial development rate of egg | 0.01098901 | day<sup>-1</sup> | (Branagan, 1973a) |
| $k_L$ | Interstadial development rate of larva | 0.03225806 | day<sup>-1</sup> | (Branagan, 1973a) |
| $k_N$ | Interstadial development rate of nymph | 0.02222222 | day<sup>-1</sup> | (Branagan, 1973a) |
| $\mu_E$ | Egg mortality rate | 0.1 | day<sup>-1</sup> | This study |
| $\mu_{LQ}$ | Mortality rate of questing larva | 0.05 | day<sup>-1</sup> | (Randolph and Rogers, 1997) |
| $\mu_{NQ}$ | Mortality rate of questing nymph | 0.03 | day<sup>-1</sup> | (Randolph and Rogers, 1997) |
| $\mu_{AQ}$ | Mortality rate of questing adult | 0.01 | day<sup>-1</sup> | (Randolph and Rogers, 1997) |
| $\mu_{LF}$ | Natural larval tick mortality | 0.005714286 | day<sup>-1</sup> | (Walker et al., 2014) |
| $\mu_{NF}$ | Natural nymphal tick mortality | 0.003703704 | day<sup>-1</sup> | (Walker et al., 2014) |
| $\mu_{AF}$ | Natural adult tick mortality rate | 0.0025 | day<sup>-1</sup> | (Walker et al., 2014) |
| $\mu_{LD}$ | Mortality rate of developing larva | 0.177388 | day<sup>-1</sup> | This study |
| $\mu_{ND}$ | Mortality rate of developing nymph | 0.065 | day<sup>-1</sup> | This study |
| $\mu_{AD}$ | Mortality rate of developing adult | 0.02 | day<sup>-1</sup> | (Randolph and Rogers, 1997) |
| $\mu_{AO}$ | Mortality rate of egg-laying female adult | 0.02 | day<sup>-1</sup> | This study |
| $\alpha_L$ | Attachment rate by larva | 0.08333333 | day<sup>-1</sup> | (Branagan, 1973a) |
| $\alpha_N$ | Attachment rate by nymph | 0.05 | day<sup>-1</sup> | (Branagan, 1973a) |
| $\alpha_A$ | Attachment rate by adult | 0.03571429 | day<sup>-1</sup> | (Branagan, 1973a) |
| $d_L$ | Feeding duration of larval tick | 5 | days | (Branagan, 1973a) |
| $d_N$ | Feeding duration of nymphal tick | 6 | days | (Branagan, 1973a) |
| $d_A$ | Feeding duration of adult tick | 8 | days | (Branagan, 1973a) |
| $P_{HN}$ | Probability nymph infects susceptible host | 0.09 | No unit | (Walker et al., 2014) |
| $P_{HA}$ | Probability adult tick infects susceptible host | 0.9 | No unit | (Walker et al., 2014) |
| $P_{LI}, P_{NI}$ | probability of a larva and nymph tick becoming infected when feeding on an acutely infectious host | 0.118656 | No unit | (Medley et al., 1993) |
| $P_{LC}, P_{NC}$ | probability of a larva and nymph tick becoming infected when feeding on an infectious carrier host | 0.023 | No unit | (Medley et al., 1993) |
| $P_{LT}$ | Probability of a feeding larva coming into contact with treatment during spray | 0.7 | No unit | This study |
| $P_{NT}$ | Probability of a feeding nymph coming into contact with treatment during spray | 0.8 | No unit | This study |
| $P_{AT}$ | Probability of a feeding adult tick coming into contact with treatment during spray | 0.9 | No unit | This study |
| $\zeta$ | Proportion females | 0.5 | No unit | This study |
| $\mu_H$ | Natural host mortality | 0.0006859604 | day<sup>-1</sup> | (Medley et al., 1993) |
| $\mu_I$ | Mortality due to East Coast fever | 0.0035 | day<sup>-1</sup> | (Medley et al., 1993) |
| $P_I$ | Probability of hosts dying from East Coast fever | 0.05 | No unit | (Medley et al., 1993) |
| $\sigma_H$ | Rate of host recovery from disease | 0.06666667 | day<sup>-1</sup> | (Medley et al., 1993) |
| $\vartheta_{IDF}$ | Mortality rate of ticks due to treatment with EPF | 9.886674 | No unit | (Oundo et al., 2024) |
| $\tau_L$ | Increased feeding duration of larva due to EPF treatment | 1.9 | days | (Nana et al., 2015) |
| $\tau_N$ | Increased feeding duration of nymph due to EPF treatment | 2.3 | days | (Nana et al., 2015) |
| $\tau_A$ | Increased feeding duration of adult tick due to EPF treatment | 3 | days | (Nana et al., 2015) |
| $k_\tau$ | Increased pre-oviposition period of the engorged female tick due to EPF treatment | 2.3 | days | (Nana et al., 2015) |
| $\tau_o$ | Increased oviposition duration | 6 | days | (Nana et al., 2015) |
| TFR | treatment fecundity reduction | 0.369 | No unit | (Nana et al., 2015) |
| $K_T$ | Environment's carrying capacity for the tick population | 35514150 | | This study |

## Results

The results shown here portray the conservative estimate of the impact of EPF. Unless explicitly stated, the default treatment strategy involves treating the cattle every two weeks for one year, and a maximum of 40% of the population receives the treatment. The duration of effective acaricidal activity of EPF is one day. The default parameter values are listed in Table 1.

### Entomopathogenic fungi marginally reduce the transmission of ECF in the cattle population

The simulation projects a slight reduction of acute infections by 11% relative to the baseline equilibrium infections after one year of treatment (Figure 2A). The EPF contributes to the observed decrease in acute infections in the cattle population by reducing the ratio of feeding ticks to cattle (Figure 2B) and limiting exposure to ticks in both treated and untreated cattle (Figures 2C and 2D), thus offering community-level protection. This is however insufficient to break the transmission cycle. Discontinuation of treatment will cause a resurgence of ECF cases in the population, an increase in the tick-to-host ratio, and an increase in cattle exposure rate to ticks.

**Figure 2.**
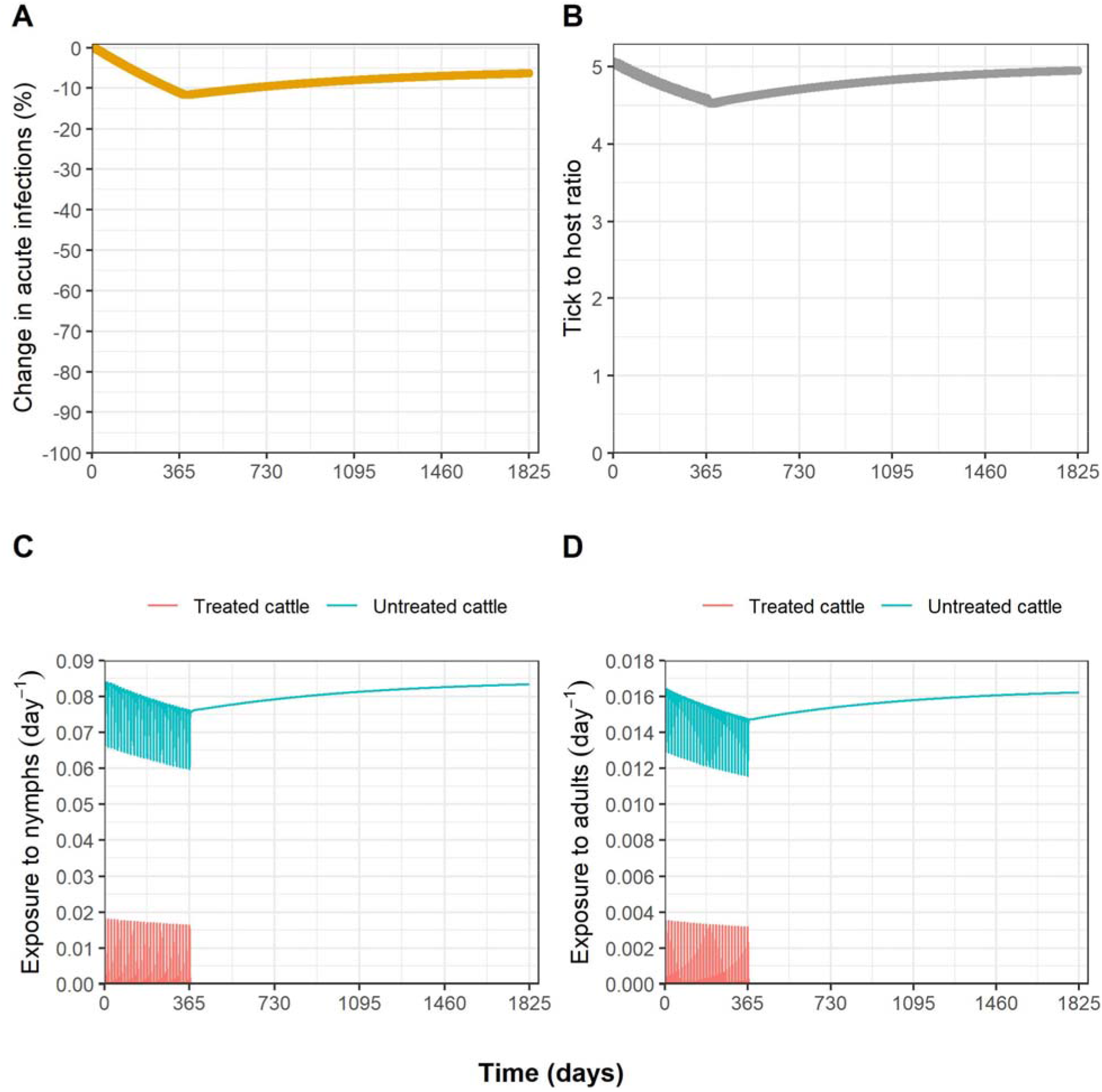
Epidemiological and entomological effects of entomopathogenic fungi. **(A)** Effect on acute *T. parva* infection in cattle population relative to the baseline equilibrium cases, **(B)** Effect on tick to host ratio, **(C)** Effect on cattle exposure to nymphal ticks, **(D)** Effect on cattle exposure to adult ticks. The treatment was applied to 40% of the host population at biweekly intervals for one year. The total simulation duration is five years and the duration of effective acaricidal activity in the EPF is one day.

### The population-level impact of entomopathogenic fungi depends on the implementation strategy

The estimated community-level impact on relative reductions of acute infections in the host population was projected to be minimal to modest depending on the coverage level and treatment frequency (Figure 3). When considering the conservative scenario of the best coverage of 40% with the biweekly treatment interval, the simulation projects a marginal decrease of 11% within one year in equilibrium acute infections. In comparison, the weekly treatment regimen with EPF emerges as the most effective, resulting in a 20.8% reduction in acute infections relative to the baseline equilibrium. Increasing the population coverage level and treatment frequency will result in a further reduction in acute infections within the cattle population. Nevertheless, none of the treatment strategies in the current product profile can cause a 50% reduction in cases of acute infections in the host population.

**Figure 3.**
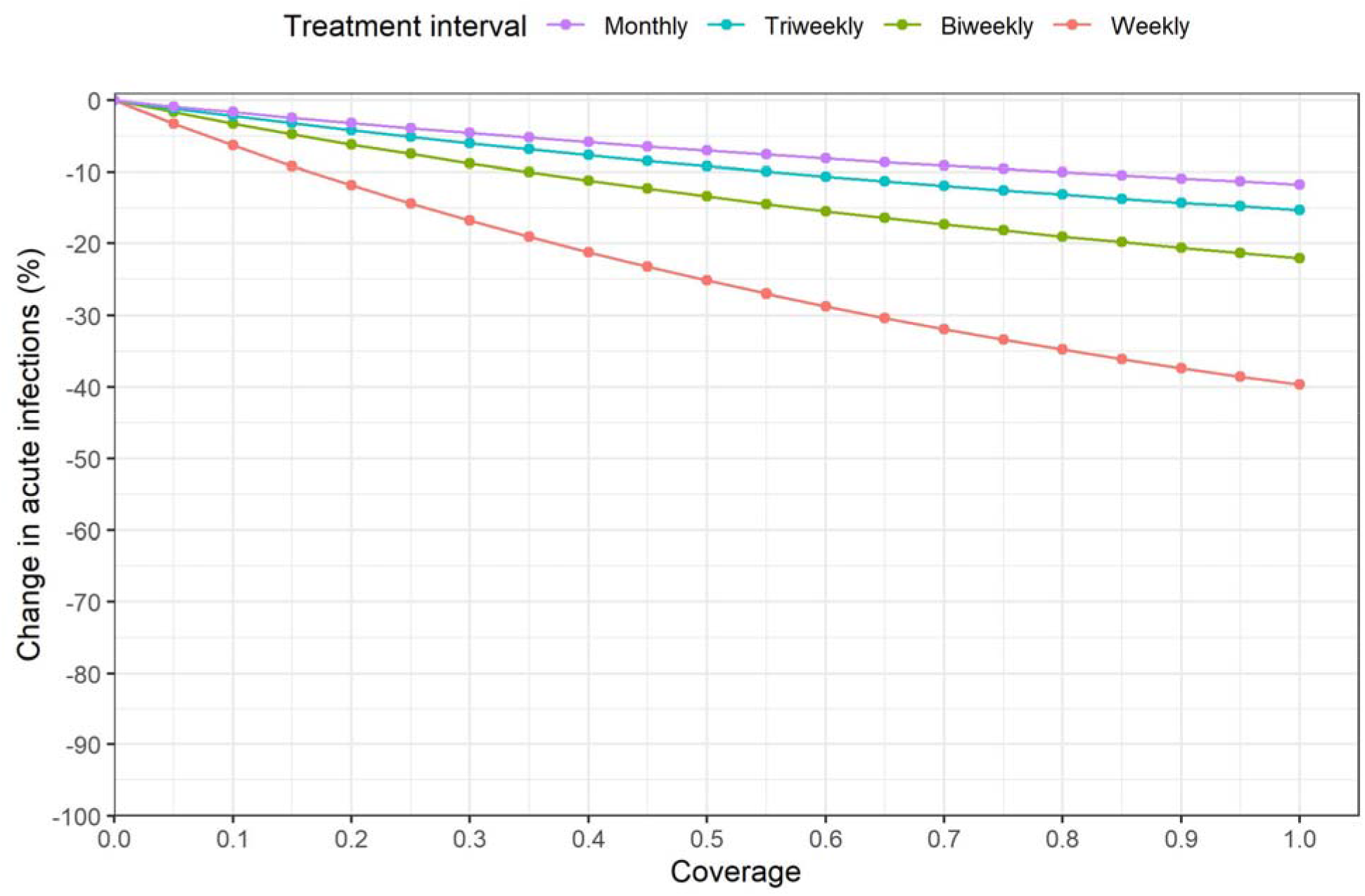
Effects of different treatment implementation strategies on the reduction of acute *T. parva* infections in the cattle population relative to the baseline equilibrium cases. Effects were assessed as a function of population coverage for different treatment intervals. The simulation period is one year and the duration of effective acaricidal activity in the EPF is one day

### Extending fungal decay time enhances the impact of entomopathogenic fungi

The model output suggests that optimizing the EPF formulation by increasing the fungal decay time will greatly improve the population-level impact of EPF, even at low coverage levels (Figure 4). For example, in the case of the most conservative estimate, which entails attaining 40% coverage through treatments administered every two weeks, and assuming a conservative estimate of effective acaricidal activity lasting one day, the simulation forecasts a slight reduction of 11% in acute infections compared to the baseline equilibrium (Figure 4B). However, extending the fungal decay time to three days leads to a 29.1% reduction, while a five-day decay period results in a 43.4% decrease. A fungal decay time of seven days causes a moderate 54.4% decline, and a ten-day decay period yields a substantial 66.4% reduction in acute infections relative to the baseline (Figure 4B). Nonetheless, the model projects that in the event it is not feasible to achieve a decay period of more than one day, then the population-level impact of EPF is maximized by high population coverage and higher treatment frequency (Figure 4A). Decreasing treatment frequency to monthly intervals will offset the impact of EPF (Figure 4C). A 50% reduction in cases of acute infections is achievable with an increase in fungal decay time.

**Figure 4.**
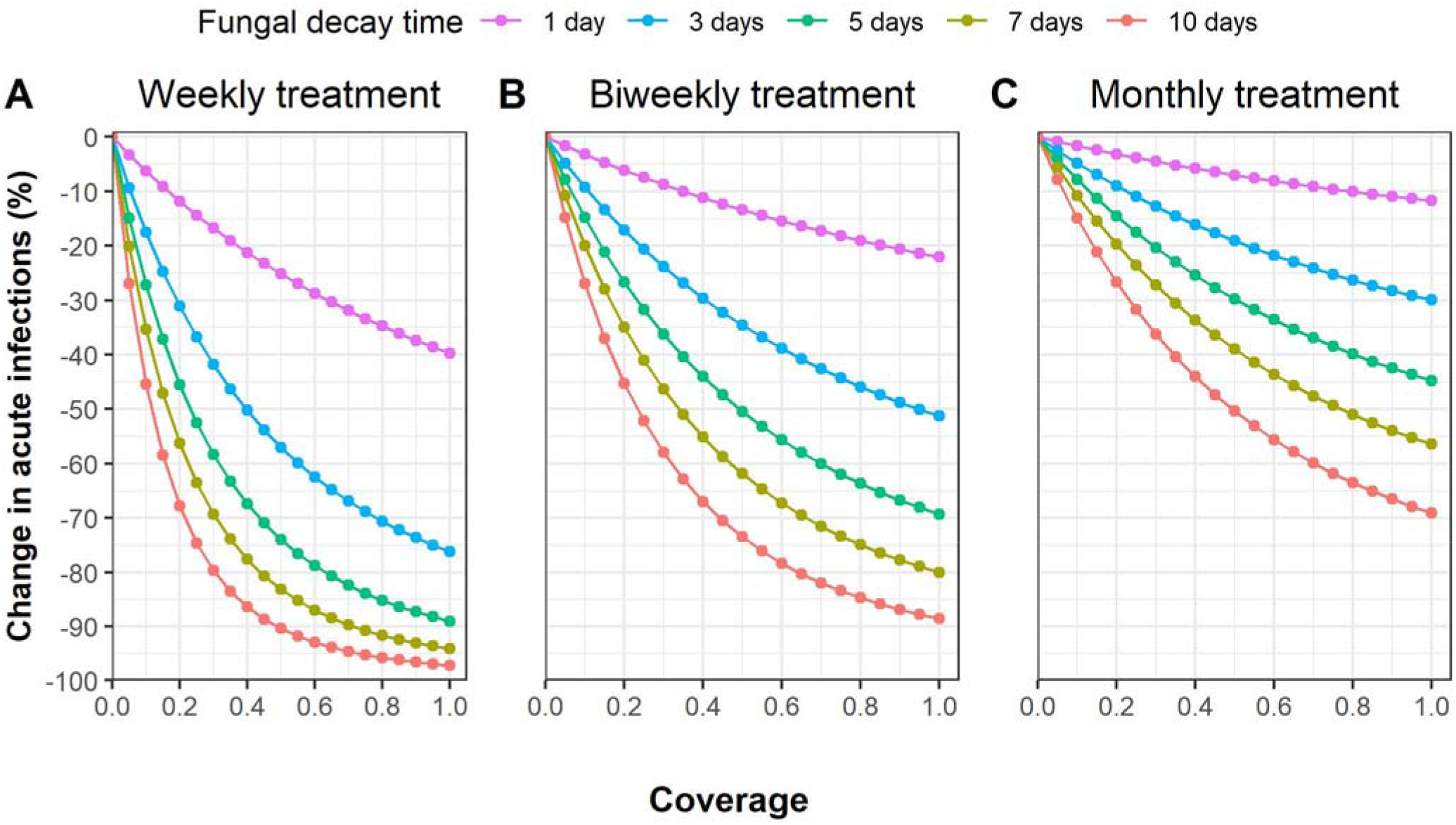
Effects of extending fungal decay time on the reduction of acute *T. parva* infections in the cattle population relative to the baseline equilibrium cases. Effects were assessed as a function of population coverage for different fungal decay times and treatment intervals. The simulation period is one year.

### The impact of entomopathogenic fungi on ECF transmission is expected to derive from the fungal mortality effect

The model simulations show that fungal-induced mortality of ticks accounts for the majority of the EPF’s impact (Figure 5). The extent of this effect will depend on the coverage and treatment frequency. In the absence of the fungal mortality effect on ticks, EPF will elicit a negligible reduction of acute infections in the cattle population (Figure 5B). This shows that the other effects of fungal infection on ticks, even when they act simultaneously, have limited potential to reduce cases of acute infections within the cattle population (Figure 5B). Notably, the model output also shows that the EPF effect of prolonging tick engorgement duration on the host is, in fact, increasing the transmission of infection in cattle, and thus exerting a minor deleterious effect on the performance of EPF (Figure 5E).

**Figure 5.**
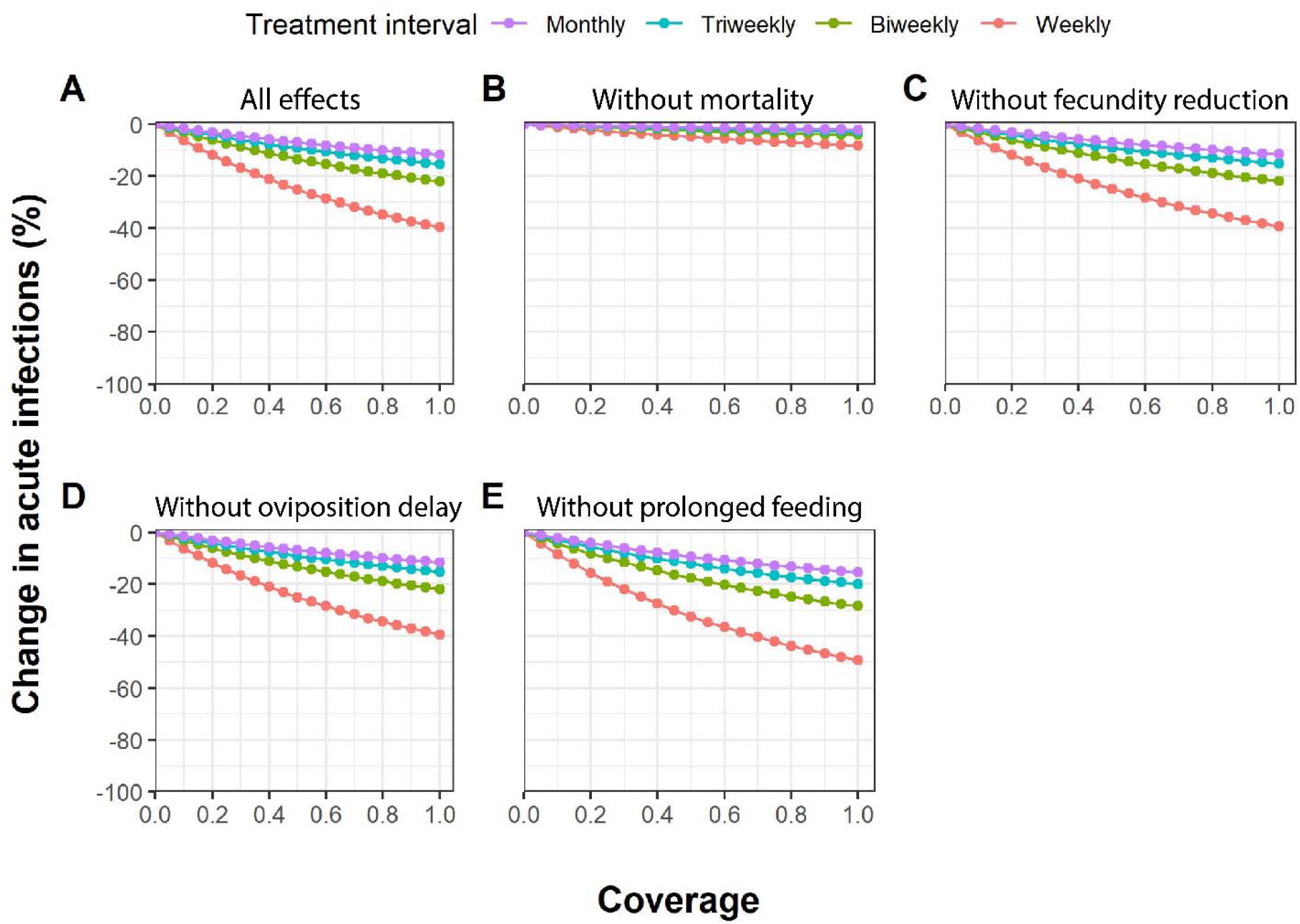
Composite effects of *Metarhizium anisopliae* ICIPE 7 product properties on the impact of entomopathogenic fungi on the acute *T. parva* infections in cattle population. Effects were assessed as a function of population coverage for different modes of action and treatment intervals. **(A)** all effects, i.e., increased mortality, reduced fecundity, delayed oviposition period, and prolonged engorgement duration, **(B)** as A without mortality effect, **(C)** as A without reduced fecundity effect, **(D)** as A without delayed oviposition effect, and **(E)** as A without prolonged engorgement effect. The treatment was applied to 40% of the host population (coverage level). The simulation period is one year and the duration of effective acaricidal activity of the EPF is one day.

### The impact of entomopathogenic fungi on ECF transmission is partly dependent on the efficiency of the spraying technique

Our simulation results indicate that enhancing the probability of tick contact with EPF, achieved through an efficient spraying technique, will result in varying degrees of effectiveness for the

EPF. This impact is contingent upon the treatment frequency and partly on the coverage level (Figure 6). At a higher treatment frequency (weekly), increasing the probability that a tick contacts the treatment will result in a considerable reduction in cases of acute infections in the host population, signifying a higher impact of EPF (Figure 6A). Conversely, a lower treatment frequency (monthly), even with a higher coverage level, will only result in a marginal reduction in the cases of acute infections in the population, signifying a lower impact of EPF (Figure 6C).

**Figure 6.**
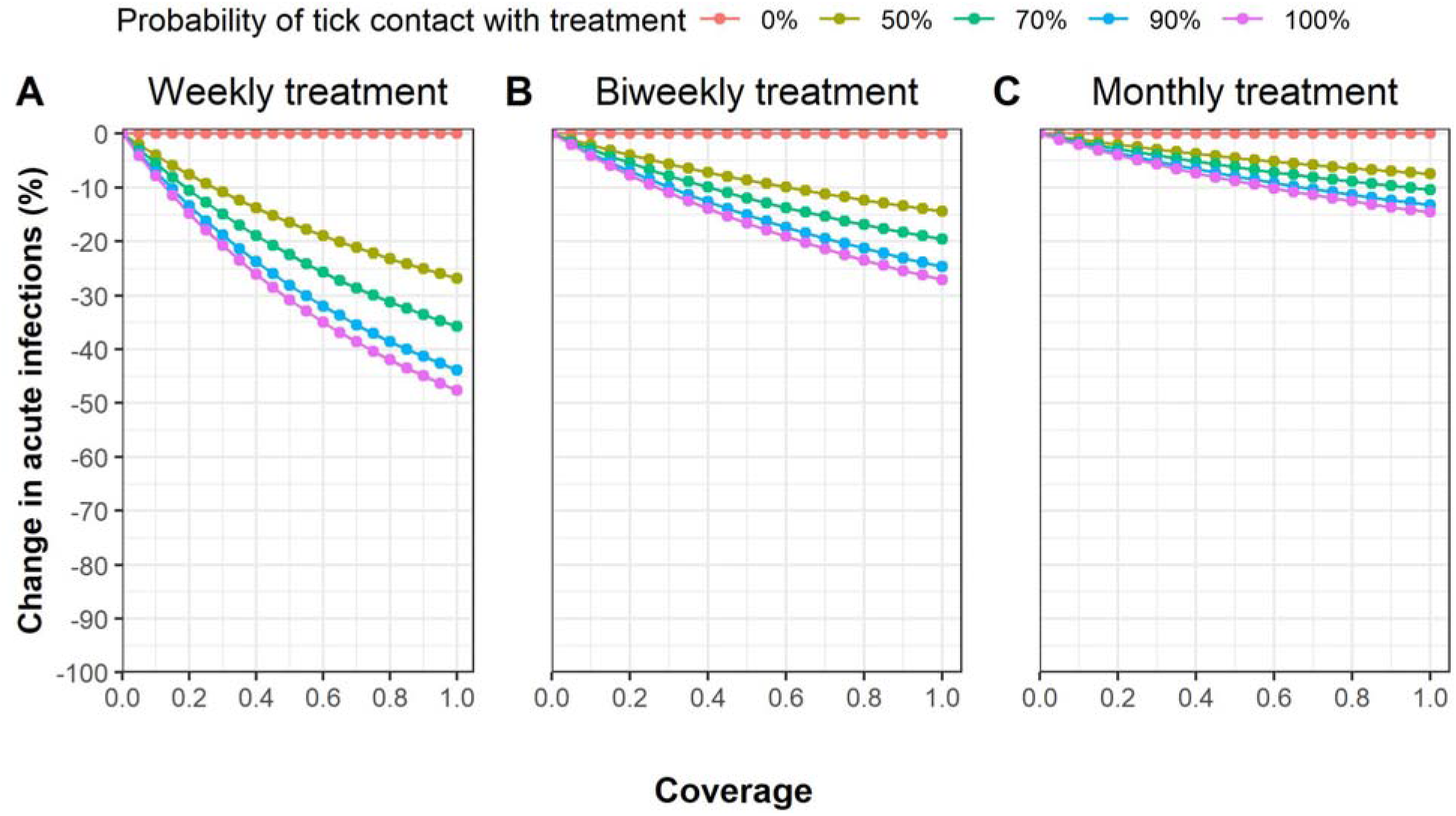
Effects of treatment efficiency and coverage level on the percentage reduction of acute infections in cattle. Effects were assessed as a function of population coverage for different treatment intervals and probabilities of tick contact with the EPF. The simulation period is one year. The probability of tick contact with EPF (*P*_*LT*_, *P*_*NT*_, *P*_*AT*_) are all the same for a given value.

## Discussion

In this study, we have developed a tick-pathogen-host interaction model to examine the potential impact of entomopathogenic fungi (EPF) on the transmission dynamics of ECF in cattle populations in an endemic context. The model was parameterized based on our results with Tickoff® biopesticide. We highlight that under the assumed product profile, EPF derives most of its impact on ECF through indirect protection: it does not prevent feeding ticks from picking up or transmitting ECF to treated animals, but it does reduce onward transmission to other animals (treated or untreated) due to delayed lethal effects. High levels of population coverage and frequent applications are needed to reduce the tick population and reach a meaningful impact on cattle populations. Substantial improvements can be obtained by improving the stability of EPFs on the cattle skin: an increase in the decay time of EPF from 1 to 7 days leads to a considerable reduction in acute infections when combined with appreciable population coverage levels, treatment intervals, and efficient spraying technique. Whether sufficient coverage levels can be reached, is also determined by the relative importance of wildlife in maintaining tick-reservoir hosts and may vary between settings.

### Mechanisms by which entomopathogenic fungi offer indirect protection against ECF in the cattle population

The rate at which susceptible cattle become infected with *T. parva* depends on several factors, including the rate of cattle exposure to host-seeking ticks, the prevalence of infection among ticks and the probability that cattle become infected after being bitten by an infectious tick. The prevalence of infection among ticks depends further on the probability that a tick becomes infected upon biting infectious cattle, the prevalence of infection in cattle, the daily tick mortality rate, and the rate of development in ticks (i.e., molting rate). Whether these factors accumulate to cause outbreaks, can be informed by the basic reproductive number, *R*_0_ (defined as the average number of newly infected cattle that arise from a single infected cattle over the course of its infectious period, in a fully susceptible cattle population). Major outbreaks may occur in naive populations only if *R*_*0*_ is greater than 1, whereas transmission will certainly die out if *R*_*0*_ is less than 1. Calculating the *R*_*0*_ for this complex system, where the epidemiological potential of a tick depends on the life stage at which it became infected, will require the construction of a next-generation matrix (NGM) model (Diekmann et al., 2010, 1990; Hartemink et al., 2008). One key component of the NGM, and part of the definition of *R*_*0*_ for this system, is the expected number of ticks feeding on a cow. A threshold for the number of ticks per cow could be calculated, above which the *R*_*0*_ would be higher than 1, and below which the transmission would stop. This threshold could give insights in how much (ongoing) treatment would be required to eradicate the pathogen, without necessarily eradicating the ticks, which may prove difficult and perhaps not even desirable as ticks have a role in (natural) ecosystems.

Whereas the rapid killing effect of synthetic acaricides can offer direct protection to treated cattle against pathogen transmission from infectious ticks, the slow-acting EPFs can only provide indirect protection to other animals (treated or untreated) by killing the infected and infectious ticks after feeding. Our model experiment indicates that EPF will indirectly reduce onward transmission of *T. parva* to other cattle by increasing the mortality rate of ticks, thus reducing the probability of surviving until the next feed and therefore transmitting the pathogen. Further, the killing of ticks by EPFs leads to reductions in the ratio of ticks to cattle and hence reduced probability of the cattle (both treated and untreated) of encountering ticks to transmit to, or acquire infection from.

Transmission of *T. parva* from *R. appendiculatus* ticks to the host animal begins at 72 hours post-tick attachment (Konnai et al., 2007). Therefore, EPF can only offer direct protection to treated cattle if it kills the infectious tick before it starts transmitting the pathogen to cattle. Although attempts have been made to enhance the virulence (i.e., the average length of time it takes to kill the tick) of EPFs through genetic manipulations (St. Leger et al., 1996; St. Leger and Wang, 2010), there is a need for further studies to investigate if these genetically modified EPFs can offer direct protection to the treated cattle against ECF.

Although laboratory experiments have demonstrated the multifaceted effects of EPFs including increased mortality, increased engorgement duration, decreased engorgement weight, and reduced reproductive fitness (Nana et al., 2015), our model simulations indicate that the impact of EPF derives most strongly from the fungus’ mortality effect. The relative contribution of this mortality effect depends on the proportion of cattle treated in the population and the frequency of re-treatment. In the absence of mortality effects, the model predicts an inconsequential impact from other modes of action.

### Product properties needed for entomopathogenic fungi to achieve a maximum epidemiological effect

Our model framework can also be used as a tool to inform what product properties are desired to obtain a better epidemiological outcome. A significant challenge in deploying fungal formulations in the field is the rapid inactivation of the conidia, and a delay in the germination process of the surviving conidia due to environmental factors such as ultraviolet (UV) radiation, low humidity, and extreme temperatures (Braga et al., 2001; Fernandes et al., 2012; Rangel et al., 2004). Conflicting findings exist regarding the average duration of persistence of *M. anisopliae* conidia on cattle skin post-treatment, ranging from up to three weeks (Kaaya et al., 1996) to up to 72 hours after application (Polar et al., 2008). Our model result shows that even at the most conservative parameter value of 40% coverage, a longer fungal decay time from 3-10 days will result in a 29.1%–66.4% reduction in the incidence of ECF infection in the cattle population compared to 11% for a decay time of 1 day. To date, techniques that have been explored for improving the persistence of EPF on treated surfaces include encapsulation of fungal conidia (Meirelles et al., 2023), incorporation of UV protectants in the formulation (Hedimbi et al., 2008), and use of thermotolerant strains of EPF (Gava et al., 2022). This may not only protect conidia from environmentally adverse conditions but also potentially increase the effectiveness of fungal formulations in natural field conditions. Further studies would be needed to ascertain the net effect of these advanced formulations of EPF.

### Implementation strategy needed to achieve a maximum epidemiological effect

Our model indicates that the projected epidemiological impact of EPF will depend on the context of their deployment strategy i.e., the treatment frequency, the population coverage level, and the efficiency of spraying. At sufficient coverage levels and treatment frequency, a large proportion of on-host *R. appendiculatus* ticks are likely to come into contact with the treatment and in doing so experience the mortality effect of EPF (Nana et al., 2015). At present, there is no standardized guideline on the optimal frequency for the application of EPFs. Previous trials have used either weekly (Murigu et al., 2016), biweekly (Alonso-Díaz et al., 2007; Oundo et al., 2024), biweekly (Alonso-Díaz et al., 2007; Oundo et al., 2024), or triweekly (Kaaya et al., 2011) treatment intervals. Our model shows an unqualified benefit of weekly (or shorter) treatment intervals for each coverage level. This strategy may, however, be logistically undesirable, especially for large herds in resource-poor settings. The biweekly treatment interval offers the best alternative.

Poor application techniques of EPF on cattle, such as improper dilution, insufficient fungal concentrations, and low spraying pressure can limit the probability of attached ticks encountering the EPF. This can further be reduced due to failure to reach the hard-to-reach predilection sites for *R. appendiculatus* such as the ear pinna, tail brush, and perianal region (Walker et al., 2003). Our model demonstrates that increasing the efficiency of the spraying technique could result in a considerable improvement of the epidemiological impact of EPFs, particularly in scenarios with high treatment frequencies. This finding of our model underscores the importance of adhering to best practices in biopesticide application, emphasizing the need to enhance spraying efficiency while maintaining a sufficient treatment frequency.

### Model limitations and future directions

Our model framework explicitly models the life cycle of ticks by incorporating the different development stages and phases, including the infection dynamics within the tick. Further, all the effects of EPF on ticks at different phases are modeled explicitly. Nevertheless, some parameter values were poorly known, especially the mortality and development rates for some immature stages. These parameters were chosen such that a population equilibrium was achieved. In our model, tick bites are assumed to be evenly divided over hosts, whereas in reality, tick burden may vary between hosts. Future studies could attempt to incorporate this exposure heterogeneity in the model.

The variations between dry and rainy seasons impact the population dynamics of *R. appendiculatus* ticks, by influencing their questing activity, development rate, survival rate, and thereby overall seasonal abundance (Randolph, 1994). Additionally, seasonal variations may affect the performance of the EPFs, with the highest efficacy occurring during the rainy season and soon thereafter (Kaaya et al., 2011; Maranga et al., 2005). Future studies could extend our model framework to incorporate such seasonality aspects.

Further, the current model is implemented in the context of a domestic environment where there is no interaction between cattle and wildlife hosts. However, *T. parva* is a multi-host pathogen with a transmission cycle involving domestic cattle and/or the Cape buffalo (*Syncerus caffer*), the wildlife reservoir host (Nene et al., 2016). The presence of this wild reservoir host may reduce the effectiveness of treatment programs, by reducing the effective coverage that can be reached. Similarly, alternative hosts such as rodents, hares, and other small mammals that may feed immature tick stages (Walker et al., 2003), and so act as tick amplifiers, are not included in the model. Their presence may, too, result in a lower chance of ticks coming into contact with EPFs. Future studies could therefore extrapolate our model framework to the wildlife-livestock interfaces.

We investigated several treatment effects of EPF on ticks. The effect of the kerosene component in the Tickoff® formulation is however not explicitly modeled due to the current lack of knowledge regarding the effects of low kerosene concentration. Nevertheless, formulations containing kerosene have elicited mortality effects on ticks (George et al., 2004; Oundo et al., 2024), sand fleas (Enwemiwe et al., 2020), and immature stages of mosquitoes (Djouaka et al., 2007; Ojianwuna and Enwemiwe, 2022). It is therefore likely that our model results are underestimating the impact of EPF formulated in kerosene.

Despite the various assumptions, our model captures essential components of the biology of tick-pathogen-host interactions relevant to the transmission dynamics of ECF, and this allowed for an extensive assessment of the epidemiological impact of EPF. The model results should however not be interpreted as predictive, but rather a demonstration of how EPF could potentially contribute to the control of ECF when deployed at a population level.

## Conclusion

The model developed here can enhance our comprehension of both the direct and indirect effects of treatments with entomopathogenic fungi, which are difficult to assess in RCTs (Reiner et al., 2016). While the model is developed for EPFs and placed in the context of the pathogen *T. parva* and *R. appendiculatus* ticks, the model can be readily adapted to other tick species, tick-borne pathogens, tick control tools, and vaccination strategies. The results from our model framework are encouraging and can be used as a basis to advocate for increased financial support towards further development of this novel tool for tick control. Further cost projections and community acceptance and uptake studies are also needed to evaluate the economic impact and cost-effectiveness of different deployment strategies.

## Abbreviations

ECF: East Coast fever
EPF: Entomopathogenic fungi
RCT: randomized controlled trial

## CRediT author statement

**Joseph Wang’ang’a Oundo:** Methodology, Formal analysis, Writing – Original Draft. **Nienke Hartemink:** Methodology, Writing – Original Draft, Writing - Review & Editing. **Mart C.M de Jong:** Conceptualization, Writing - Review & Editing. **Constantianus J.M. Koenraadt:** Writing - Review & Editing. **Shewit Kalayou:** Writing – Review & Editing. **Daniel Masiga:** Conceptualization, Writing - Review & Editing, Funding acquisition. **Quirine ten Bosch:** Conceptualization, Methodology, Writing – Original Draft, Writing - Review & Editing.

## Acknowledgments

We are grateful to Mariken de Wit and You Chang of the Quantitative Veterinary Epidemiology group who offered support during the coding of the modeling script in R.

## Availability of data and materials

The code used to perform the model analyses will be available on GitHub.

## Consent for publication

Not applicable

## Appendix A. Supplementary data

## Appendix A.1: Variables in the model

| Variable | Description |
| --- | --- |
| $E_{US}$ | Oviposited eggs from female tick |
| $LQ_{US}$ | Untreated and susceptible questing larva |
| $LF_{US}$ | Untreated and susceptible feeding larva |
| $LD_{US}$ | Untreated and susceptible developing larva |
| $LD_{TS}$ | Treated and susceptible developing larva |
| $LD_{UI}$ | Untreated and infected developing larva |
| $LD_{TI}$ | Treated and infected developing larva |
| $NQ_{US}$ | Untreated and susceptible questing nymph |
| $NQ_{UI}$ | Untreated and infectious questing nymph |
| $NF_{US}$ | Untreated and susceptible feeding nymph |
| $NF_{UI}$ | Untreated and infectious feeding nymph |
| $ND_{US}$ | Untreated and susceptible developing nymph |
| $ND_{TS}$ | Treated and susceptible developing nymph |
| $ND_{UI}$ | Untreated and infectious (and newly infected) developing nymph |
| $ND_{TI}$ | Treated and infectious (and newly infected) developing nymph |
| $AQ_{US}$ | Untreated and susceptible questing adult |
| $AQ_{UI}$ | Untreated and infectious questing adult |
| $AF_{US}$ | Untreated and susceptible feeding adult |
| $AF_{UI}$ | Untreated and infectious feeding adult |
| $AD_U$ | Untreated and developing adults (both male and female) |
| $AD_T$ | Treated and developing adults (both male and female) |
| $AO_U$ | Untreated egg-laying adult females |
| $AO_T$ | Treated egg-laying adult females |
| $H_{US}$ | Untreated and susceptible cattle host |
| $H_{UI}$ | Untreated and infectious cattle host |
| $H_{UC}$ | Untreated and carrier cattle host |
| $H_{TS}$ | Treated and susceptible cattle host |
| $H_{TI}$ | Treated and infectious cattle host |
| $H_{TC}$ | Treated and carrier cattle host |

## Appendix A.2: Model equations

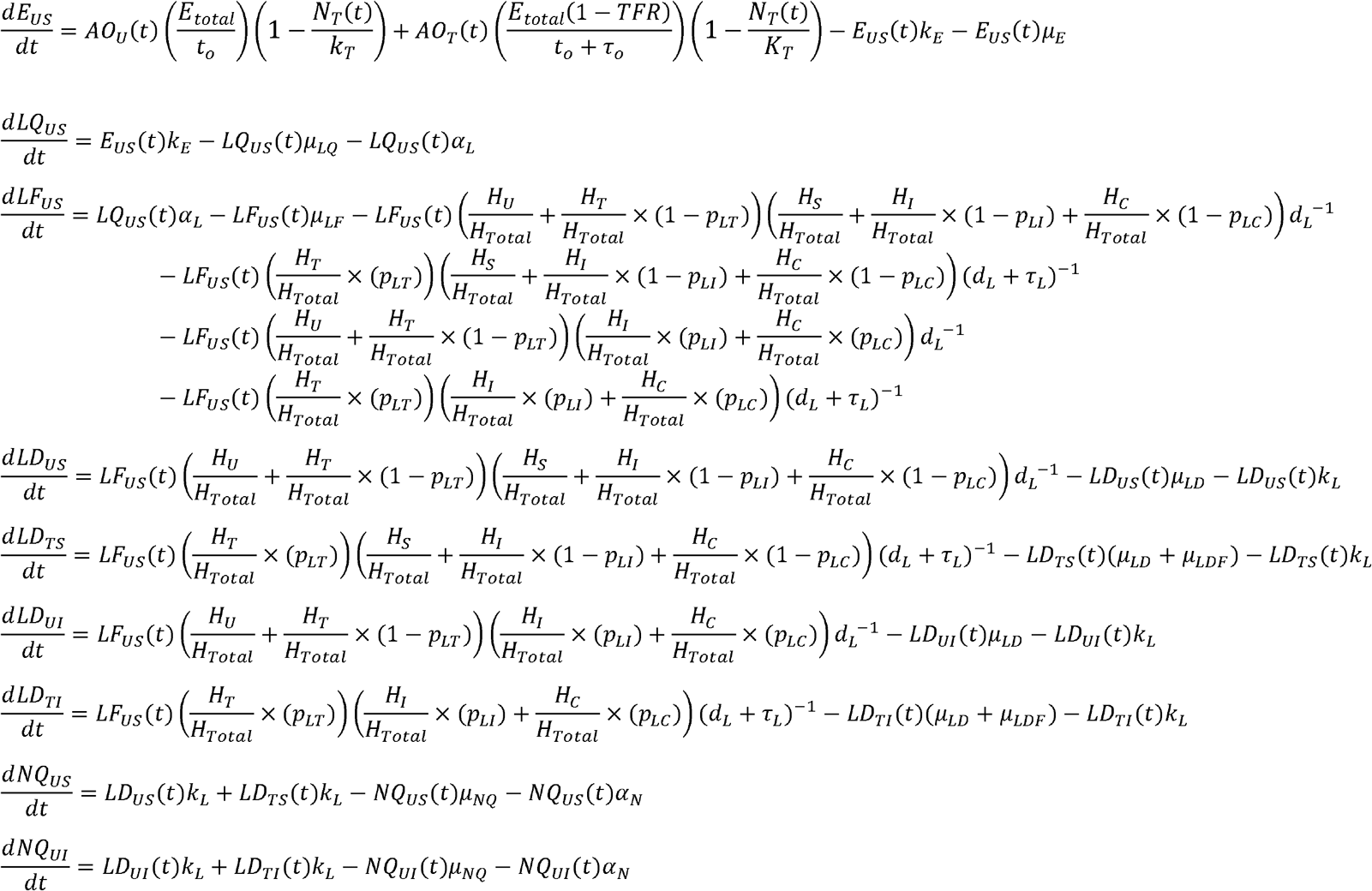

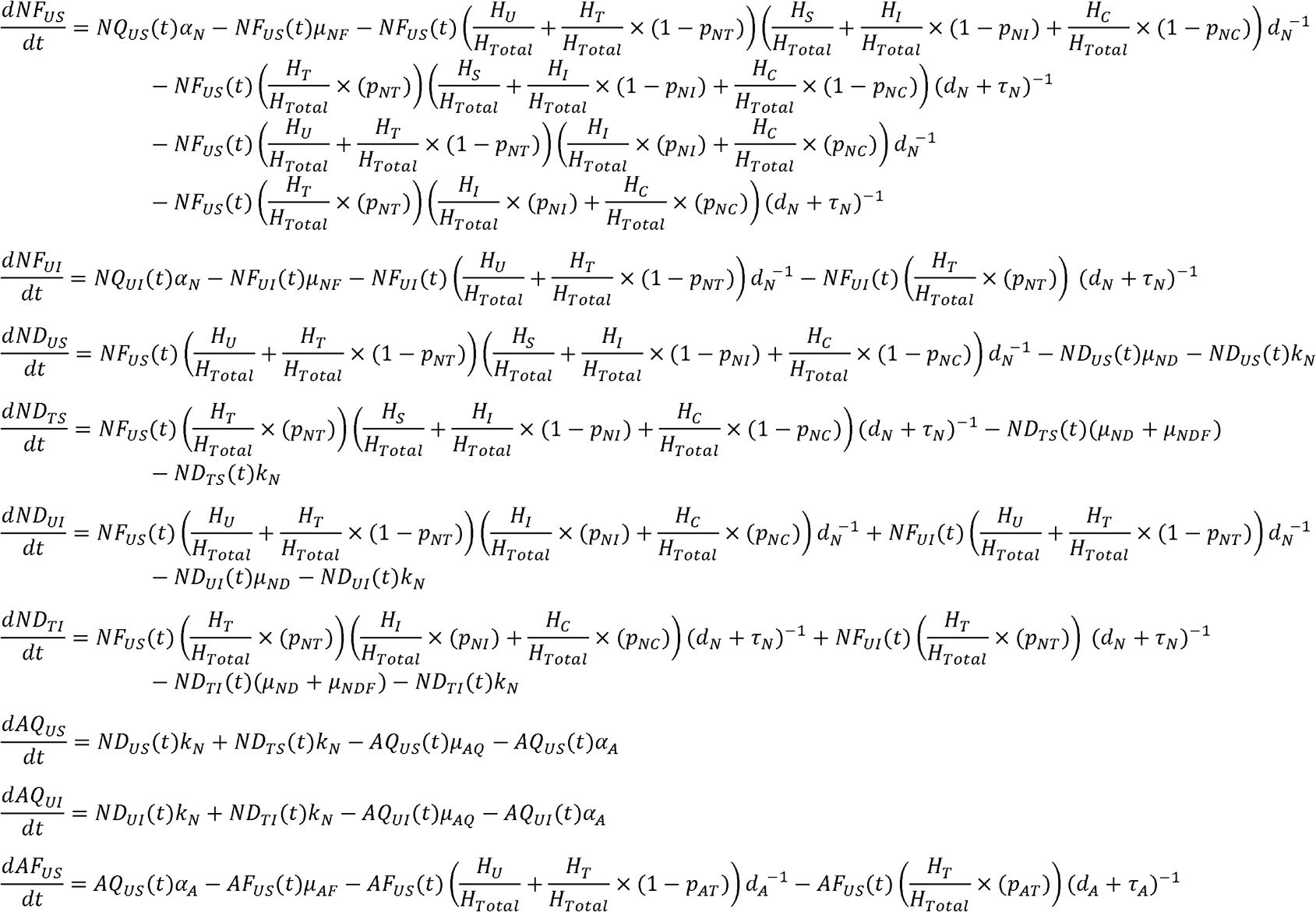

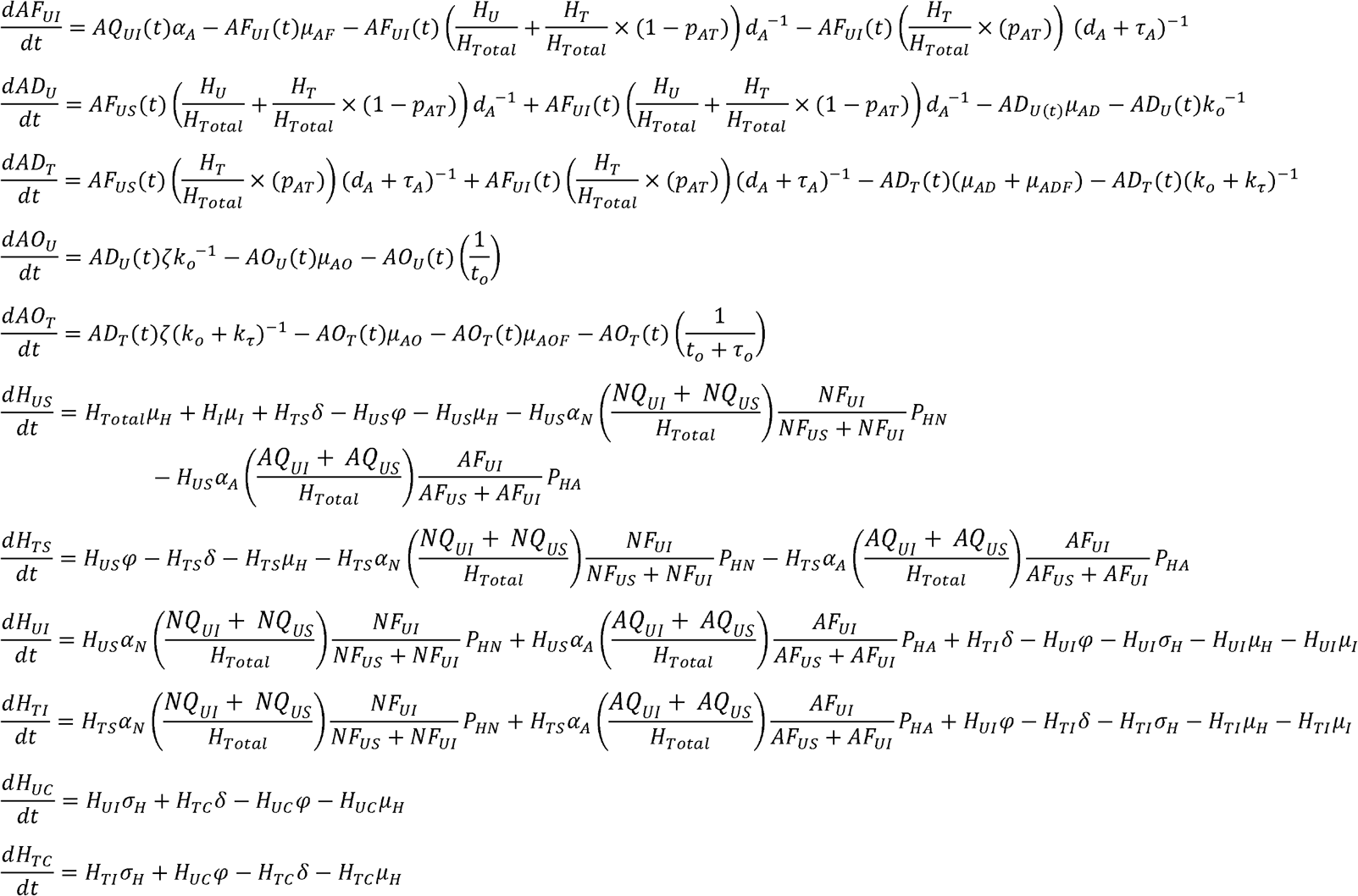

